# Super-resolution ultrasound imaging of cerebrovascular impairment in a mouse model of Alzheimer’s disease

**DOI:** 10.1101/2022.10.05.511008

**Authors:** Matthew R. Lowerison, Nathiya Vaithiyalingam Chandra Sekaran, Zhijie Dong, Xi Chen, Qi You, Daniel A. Llano, Pengfei Song

## Abstract

Increasing evidence has suggested a link between cerebrovascular disease and the cognitive impairment of patients with Alzheimer’s disease. However, cerebrovascular disease and Alzheimer’s disease share several risk factors making it unclear whether cerebrovascular deficiency and Alzheimer’s disease pathology have additive effects on cognition or if cerebrovascular impairment merely exacerbates existing Alzheimer’s disease-associated cognitive decline. Additionally, early-stage Alzheimer’s disease typically involves hippocampal atrophy, complicating most efforts to elucidate the interplay between cerebral microvascular function and Alzheimer’s disease progression due to the necessity of probing deep-brain structures. The purpose of this study was to demonstrate the use of ultrasound localization microscopy on the 5xFAD mouse model of Alzheimer’s disease (3-month and 6-month-old cohorts) in comparison to age-matched wild-type controls, revealing microvascular scale reconstructions throughout the whole brain depth, to visualize and quantify Alzheimer’s disease-associated vascular impairments. We found that functional decreases in hippocampal and entorhinal flow velocity preceded structural derangements in regional vascular density. In addition to providing global vascular quantifications of deep brain structures with a high local resolution, this technology also permitted hierarchical analysis of individual vessels and, in some cases, potentially allowed for decoupling of arteriole and venous flow contributions. Co-registered histological sectioning confirmed the regionalized hypo-perfusion deficits seen on ultrasound imaging, which were co-localized with amyloid beta plaque deposition.

**Significance statement:** The study of the impact of cerebrovascular disease on Alzheimer’s disease pathology is complicated by the need to image deep-brain structures with high vascular fidelity. We demonstrate that ultrasound localization microscopy, a super-resolution acoustic imaging technique, is capable of imaging cerebrovasculature throughout the entire depth of the brain at a microvascular scale. This technology was applied to the 5xFAD mouse model of Alzheimer’s disease, where it was found that 5xFAD animals have significant impairments in vascular function in the entorhinal cortex and hippocampal region in comparison to age matched controls at the 3-month timepoint. Structural derangements in cerebrovasculature were only observed in the 6-month-old animal cohorts, with a maintained impairment in vascular function.

## Introduction

Alzheimer’s disease (AD) is the leading cause of dementia among older adults. Approximately 6 million Americans are impacted by this disease, and this number is projected to more than double to 14 million by 2060 (Matthews et al., 2019), leading to the loss of quality of life to a substantial and growing population of individuals. Unfortunately, there are currently no effective preventative or therapeutic strategies to combat this disease, signaling an urgent need to improve early diagnosis and therapeutic options to help prevent, treat, and alleviate AD (HHS, 2021).

The current hallmarks of AD are amyloid beta (Aβ) deposition and abnormal accumulations of neurofibrillary tau protein (Bloom, 2014), motivating substantial research effort centered on the amyloid hypothesis (Glenner and Wong, 1984; Hardy and Allsop, 1991; Karran et al., 2011) of AD. However, additional pathophysiological processes likely contribute to AD. Increasing evidence has suggested a link between cerebrovascular disease (CVD) and the cognitive impairment and other symptoms of patients with AD (de la Torre, 1994), with vascular dysfunction identified as an early and nearly ubiquitous feature of AD (Iadecola, 2004; Kisler et al., 2017). However, CVD and AD share several risk factors, such as hypertension (Bellew et al., 2004), diabetes (Ahtiluoto et al., 2010), and obesity (Kivipelto et al., 2005), making it unclear whether cerebrovascular deficiency and AD pathology have additive effects on cognitive decline or if CVD merely exacerbates existing AD-associated cognitive impairments. The vascular hypothesis of AD (de la Torre and Mussivand, 1993), details a vicious cycle between vascular dysfunction and AD pathology: vascular dysregulation initiates a cascade of neurovascular dysfunction and cerebrovascular blood flow reduction leading to the accumulation of Aβ, which in turn aggravates further vascular dysregulation by triggering hypoperfusion, vasoconstriction, and breakdown of the blood-brain barrier (Kisler et al., 2017). These cerebrovascular impairments worsen Aβ clearance, further accelerating AD pathology. Understanding and overcoming this vicious cycle is essential for developing effective AD therapies and interventions.

Despite CVDs profound impact on AD development and progression, the exact mechanisms and relationships underlying a vascular hypothesis of AD remains difficult to establish due to the requirements of high-fidelity and dynamic characterization of microvascular impairments. Furthermore, early-stage AD typically involves hippocampal atrophy (Schuff et al., 2009), complicating most efforts to elucidate the interplay between cerebral microvascular function and AD progression due to the necessity of probing deep-brain structures at a resolution sufficient to image microvasculature – an intractable set of requirements for most medical imaging modalities. Histological analysis of AD-affected brain sections can give insight into the structural correspondence between microvascular density and Aβ deposition but lacks critical information on impairments in blood flow dynamics which likely precede vascular remodeling. Ultrasound localization microscopy (ULM) is an emerging super-resolution ultrasound technique (Christensen-Jeffries et al., 2015; Errico et al., 2015) that can potentially provide insight into the interplay between microvascular pathology and AD progression. ULM provides super-resolved images of patent vasculature by localizing the trajectories of microbubble contrast agents at a sub-diffraction precision, without sacrificing ultrasound imaging penetration depth. ULM has been successfully applied across a variety of organ types, including cerebrovascular vessel reconstruction in mice (Demeulenaere et al., 2022; Lowerison et al., 2022), rats (Errico et al., 2015), and in humans (Demené et al., 2021). Notably, the theoretical resolution limit of ULM approaches the capillary-scale (Desailly et al., 2015), which may permit the *in vivo* characterization of focally-constricted capillaries and microvessels that have been reported in the brains of AD patients and in AD rodent models (Nortley et al., 2019).

In this manuscript, we apply ULM to a 5xFAD mouse model of AD with age and sex-matched controls from two different stages of AD progression (3-month and 6-month-old cohorts). We examine global hippocampal, entorhinal, and isocortex alterations in the microvascular density and median blood flow velocity from these two cohorts of animals. The resulting super-resolved whole-brain cross-sectional images are then compared to co-registered histological sections of vascular staining and Aβ deposition. A case study examining velocity profiles of individual vessels in the cortex and hippocampus is also explored.

## Materials and Methods

### Animal model and ethics

All procedures conducted on mice presented in this manuscript were approved by the Institutional Animal Care and Use Committee (IACUC) at the University of Illinois Urbana-Champaign (protocol # 22033). All experiments were performed in accordance with these IACUC guidelines. Mice were housed in an animal care facility approved by the Association for Assessment and Accreditation of Laboratory Animal Care. Every attempt was made to minimize the number of animals used and to reduce suffering at all stages of the study. Reporting in this manuscript follows the recommendations of the ARRIVE guidelines (Sert et al., 2020).

5XFAD transgenic mice overexpress mutant human amyloid beta (A4) precursor protein 695 (APP) with the Swedish (K670N, M671L), Florida (I716V), and London (V717I) Familial Alzheimer’s Disease (FAD) mutations along with human presenilin 1 (PS1) harboring two FAD mutations, M146L and L286V. 5xFAD mice (MMRRC Strain # **034840-JAX**, Jackson laboratory) were crossbred and generated with B6FJLF1/J (Stock No: #100012, Jackson laboratory) mice. Transgenic Genotyping was confirmed using Transnetyxservice. Both 5xFAD and control littermates of either sex, aged 3- and 6-months old, were used in this study. Littermate control animals of same age were used to avoid confounding effects. Animal cohort sizes were 17 in the 3-month group (5xFAD = 8, WT = 9), and 24 in the 6-month group (5xFAD = 13; WT = 11), with datasets excluded if the animal died prematurely during imaging.

### Craniotomy preparation for ultrasound imaging

Mouse anesthesia was induced using a gas induction chamber supplied with 4% isoflurane mixed with medical oxygen. Mice were then placed in a stereotaxic frame with nose cone supplying 2% isoflurane with oxygen for maintenance, with 1% lidocaine intradermally injected into the scalp to supplement anesthesia. The mouse’s head was secured to the stereotaxic imaging stage via ear bars (**Figure 1A**). The scalp of the mouse was removed with surgical scissors, and a cranial window was opened on the left side of the skull using a rotary Dremel tool. The cranial window started at the sagittal suture and extended laterally to expose the lateral expanse of the cerebral cortex. Electrocardiogram (ECG) signal was acquired using an iWorx (Dover, NH, USA) IA-100B single channel biopotential amplifier with C-MXLR-PN3 platinum needle electrodes inserted into the legs of the animal to monitor heart rate.

**Figure 1.**
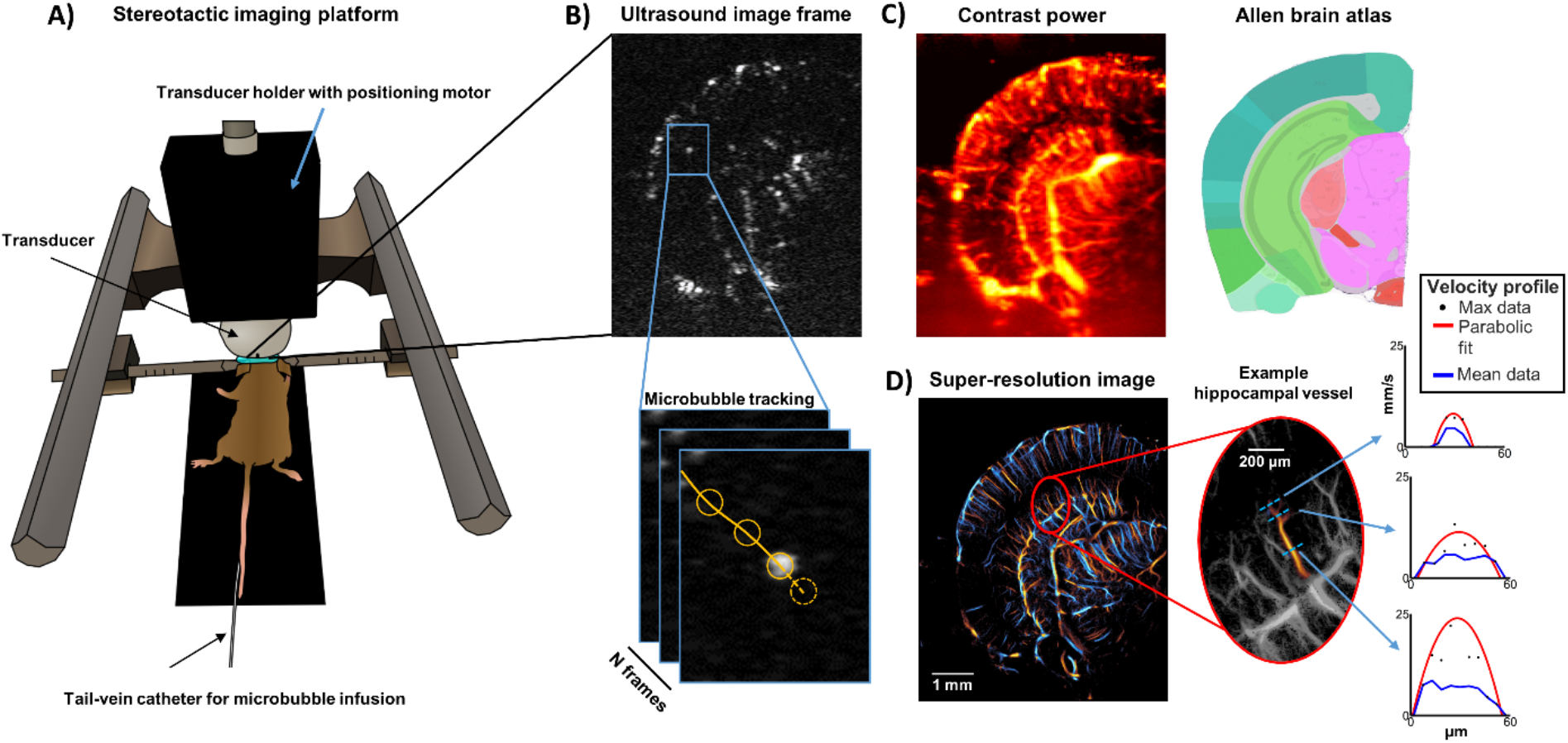
Overview of ULM imaging acquisition. **(A)** Post-craniotomy animals were secured to a stereotaxic imaging frame via ear-bars, and an ultrasound transducer with high-precision motor was positioned to acquire the anatomical imaging plane of interest. **(B)** An example imaging frame of microbubble contrast-enhanced data taken from a coronel cross-section of the brain. Individual microbubble centroids were localized and tracked to produce blood flow trajectories (orange line). **(C)** A conventional contrast-enhanced power Doppler image of the mouse brain, with corresponding Allen brain atlas to identify anatomy. **(D)** A super-resolved directional flow ULM image of the same dataset as in (C), demonstrating finer detail of the cerebral microvasculature. Individual vessels (e.g.: hippocampal vessel in red circle ROI) can be extracted and investigated to perform local analysis of blood flow velocity profiles.

### Contrast-enhanced ultrasound imaging of mouse brain

A Verasonics Vantage 256 research ultrasound system (Verasonics Inc., Kirkland, WA) was used for all ultrasound imaging in this study. A L35-16vX transducer (Verasonics) was secured to the stereotaxic imaging frame via a 3D-printed transducer holder (**Figure 1A**) with attached high-precision translation motor (VT-80 linear stage, Physik Instrumente, Auburn, MA). The transducer was oriented to produce a coronal anatomical section of the brain (**Figure 1B**). Acoustic contact gel was applied directly to the surface of the mouse brain. Once acoustic coupling was established, the motorized stage was adjusted by 0.1mm increments to find an imaging plane which contained a cross-section of hippocampus (**Figure 1C**). This was generally 3.0-3.5mm caudal to bregma. The imaging field of view was then adjusted laterally/axially to cover the entire half of the mouse brain in this anatomical position. Two additional imaging planes, one 0.5mm caudal to this center position and one 0.5mm rostral to it, were also selected for imaging to provide a more global sampling of hippocampal vasculature.

The tail vein of the mouse was cannulated with a 30-gauge catheter and vessel patency was confirmed with a 0.1 mL injection of sterile saline. Freshly activated DEFINITY® was perfused via the tail vein catheter at a rate of 10 μL/min using a programmable syringe pump (NE-300, New Era Pump Systems Inc., Farmingdale, NY). The MB solution was mixed every 5 minutes using a magnetic stirrer to maintain a constant MB concentration during the experiment. Ultrasound imaging was performed at a center frequency of 20 MHz, using 9-angle plane wave compounding (1-degree increments) with a post-compounding frame rate of 1,000 Hz (**Figure 1B**). A total of 64 seconds of data (64,000 frames) were acquired for each imaging plane. Ultrasound data was saved as beam-formed in-phase quadrature (IQ) datasets for off-line processing in MATLAB (The MathWorks, Natick, MA; version R2019a).

### Reconstruction of super-resolution brain vascular images

Singular value decomposition (SVD) clutter filtering was applied to extract microbubble signals from tissue background from each IQ dataset (Desailly et al., 2017; Lowerison et al., 2020b; Huang et al., 2020; Song et al., 2018b), with the singular value threshold determined adaptively (Song et al., 2017a). A noise-equalization profile (Song et al., 2017b) was applied to equalize the microbubble signal intensity through the entire depth of imaging. A 2D Gaussian function was empirically fit to the axial and lateral dimensions of manually selected microbubbles to represent the point-spread function (PSF) of the system. Microbubble separation (Huang et al., 2020) was applied to post-SVD-filtered IQ data, and then each IQ data subset was spatially interpolated to an isotropic 4.928 μm axial/lateral pixel dimension using 2D spline interpolation (Song et al., 2018a). Normalized 2D cross-correlation was performed on this interpolated dataset with the empirical PSF to identify candidate microbubble localizations (**Figure 1B**). Pixels with a low cross-correlation coefficient were excluded via a threshold (Huang et al., 2020; Lowerison et al., 2020b; Tang et al., 2020), and microbubble centroids were localized with the MATLAB built-in “imregionalmax.m” function. Frame-to-frame microbubble centroid pairing and trajectory estimation was performed using the uTrack algorithm (Jaqaman et al., 2008). A minimum microbubble trajectory length of 20 frames (i.e., 20 ms) was applied to the super-resolution reconstructions presented in this study. Each acquisition was then accumulated to produce a final ULM vascular reconstruction of the brain vasculature (**Figure 1D**).

### Ultrasound image analysis

Anatomical regions (**Figure 2**) were manually segmented by placing Bezier control vertices on the border of the region of interest (ROI), informed by the Allen brain atlas, and by interpolating between each vertex using Hobby’s algorithm (Hobby, 1986). Three anatomical regions of interest were selected: the isocortex, the entorhinal cortex, and the hippocampal region. Brain vascularity within each ROI was calculated by binarization of super-resolution vessel maps to determine the percentage of cross-sectional area that contained microbubbles. Blood vessel velocity was determined for every microbubble track directly from the frame-to-frame displacement of detected MB centroids. Vascular tortuosity, as measured by the sum of angles metric (SOAM), was calculated using the algorithm described by Shelton *et al*. (Shelton et al., 2015) for every MB trajectory and averaged temporally across all data accumulations. Blood velocity profiles were determined by manually selecting vessel cross-sections on the final ULM accumulation, and then extracting all raw velocity trajectories that crossed the line segmentation (**Figure 1D, Figure 3**). Parabolic profiles were fit to the raw maximum velocity estimates using MATLAB.

**Figure 2.**
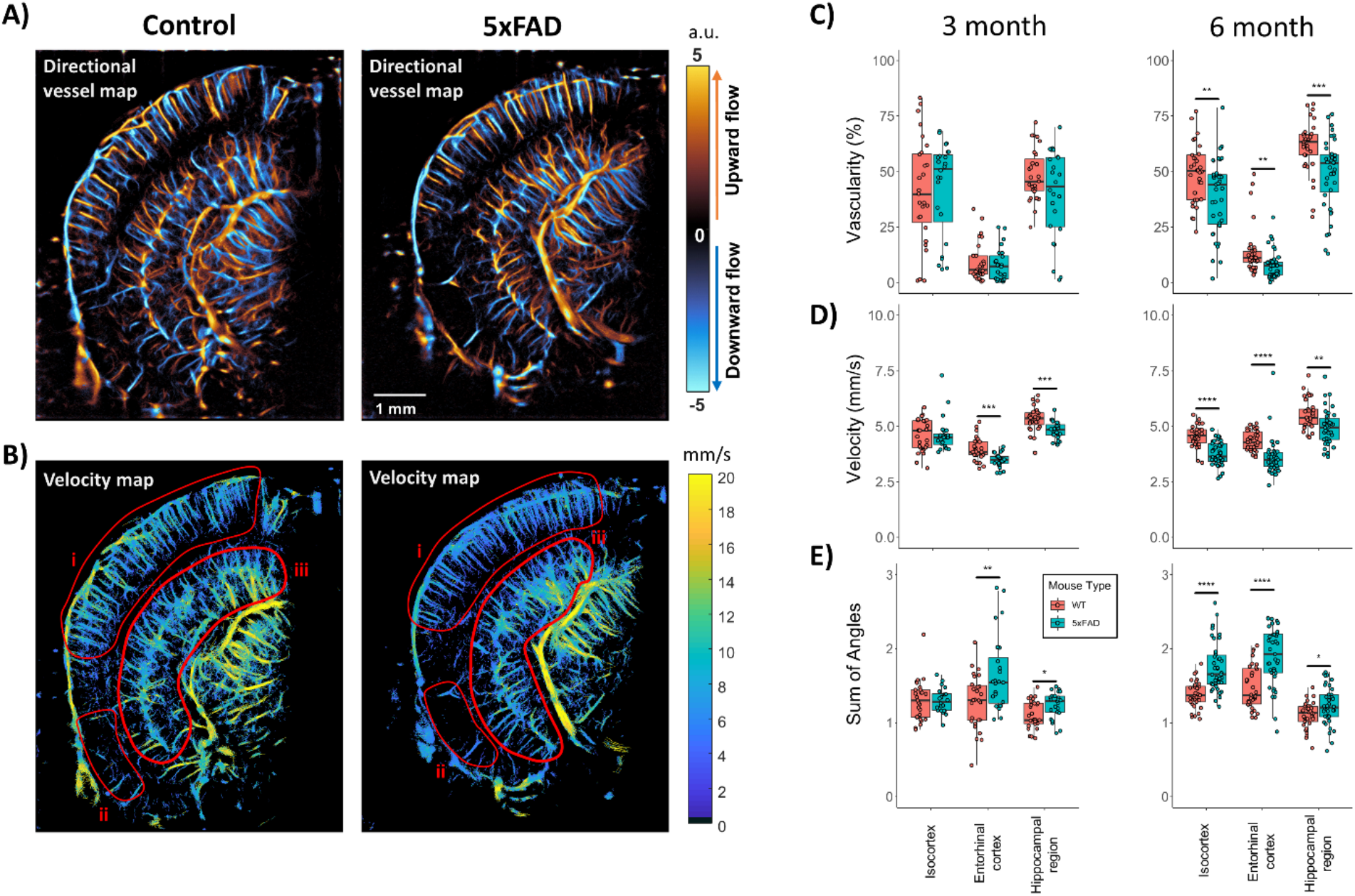
Regional anatomy analysis with ULM. **(A)** Example directional flow and **(B)** velocity maps taken from a WT control (left) and 5xFAD (right) mice in this study. Segmentations were placed around (i) dorsal isocortex, (ii) entorhinal cortex, and (iii) hippocampal region ROIs for analysis, as demonstrated on the velocity maps. **(C)** The 6-month-old 5xFAD cohort demonstrated significant decreases in vascularity across all three ROIs in comparison to WT controls. No decrease in vascularity was noted for the 3-month cohort. **(D)** A significant decrease in overall median velocity was noted in the entorhinal cortex and hippocampal region for the 3-month 5xFAD cohort, with a trend toward a decrease in the isocortex. This decrease in velocity was sustained in the 6-month-old 5xFAD cohort, with significance found in all three ROIs. **(E)** The 5xFAD cohorts had increased tortuosity, as measured by the sum-of-angles index, across all regions in the 6-month cohort, with evidence of early increased tortuosity in the entorhinal and hippocampus of the 3-month animals. (* p < 0.05, ** p < 0.01, *** p < 0.001, **** p < 0.0001).

**Figure 3.**
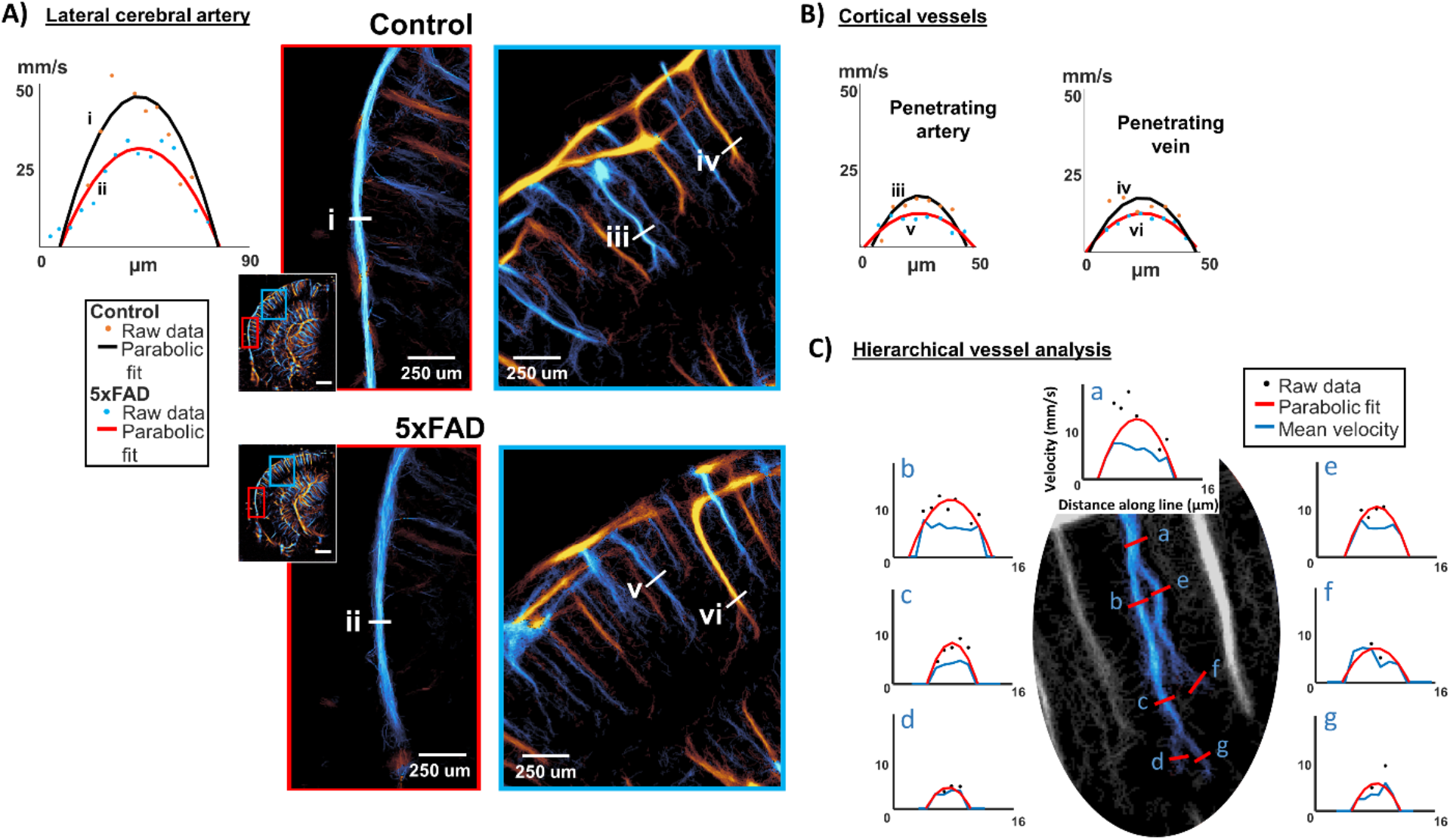
Local vessel analysis demonstrates lateral cerebral artery and penetrating cortical vessel velocity reduction. **(A)** Local ROIs from both WT control (top) and 5xFAD (bottom) animals are shown for a section of the lateral cerebral artery and for a cortical region. The velocity profiles of the lateral cerebral artery, including both raw data and a parabolic (laminar) fit, demonstrate a decrease in peak velocity for the (ii) 5xFAD animal in comparison to the (i) WT control. **(B)** A similar trend is also noted for penetrating arteries (vessel profiles iii and v) and veins (vessel profiles iv and vi) in the cortical regions. **(C)** A hierarchical analysis of a penetrating cortical artery demonstrates a decrease in peak flow velocity as the length of the vessel is traversed (from a to g), along with less evidence of pulsatility in blood flow.

### FITC dextran perfusion and tissue processing

Following the ULM imaging FITC-dextran (70,000 Kda; 0.1 mL 50 mg/mL in double-distilled water; Sigma, St. Louis, MO, USA) was administered intravenously through tail vein of mouse. After five minutes of injecting FITC-dextran, animals were sacrificed. Brains were rapidly removed and placed in 4 % paraformaldehyde (PFA) in PBS at 4 °C for 24h −48h and then cryoprotected in ascending series of 10%, 20%, 30% sucrose in PBS for another 24–36h at 4°C. Each brain was embedded and cut into 40-μm-thick sections on a cooling microtome.

### Immunostaining

Sections were washed in TBS for 3 times for ten minutes each. Sections were then incubated for 30 min in a solution of 0.3% Triton X-100 in TBS to enhance membrane permeability. The sections were then transferred to a blocking solution consisting of 0.3% Triton X-100 and 5% Normal goat serum and 3% BSA in TBS and incubated for 60 min. The primary antibody solution consisted of 1:1000 Purified anti-β-Amyloid, 1-16 Antibody (catalog# SIG-39320, Biolegend) in blocking buffer. Sections were incubated in this solution overnight and rinsed in three changes of the Triton X-100 solution the following day. The sections were then transferred to a secondary antibody solution and incubated at room temperature for 2h. This solution consisted of 1:100 Alexa Fluor 568-conjugated goat anti-mouse secondary antibody (catalog #A-11004, Invitrogen). Following a final series of washes in TBS, the sections were mounted on gelatin-coated slides and coverslipped with an anti-fade solution (Vectashield; Vector Laboratories).

### Histology imaging and analysis

Sections used for histochemical analysis were imaged using a Confocal - Zeiss LSM 710 - Multiphoton Microscope and Zen blue software. For each hippocampus tissue section, 40× mosaic Z-stacks were taken throughout the entire depth and x–y plane of the brain. The stacks were collapsed into 2D maximum intensity projections and tiled into a single image using Zen blue software.

### Experimental design and statistical analyses

All statistical analysis was performed in the R programming language (R Core Team, 2019), and all graphs were generated using the ggplot2 package (Wickham, 2016). A two-way analysis of variance (ANOVA) was applied to test for statistical significance between WT and 5xFAD mouse brain anatomy with a Tukey’s honestly significant difference test applied as a post-test. A corrected p < 0.05 was considered as statistically significant. All summary statistics in text are presented as the mean ± standard deviation.

## Results

### Super-resolution ULM provides microvascular fidelity over whole brain depth

A diagrammatic example of the imaging platform and protocol used in this experiment is demonstrated in **Figure 1A**. After craniotomy, the anesthetized mouse was secured via ear-bars to a stereotactic imaging frame with connected ultrasound transducer and high precision motor for imaging cross-section placement. After reaching a steady-state concentration, the microbubble contrast agent in the mouse brain was visible as distinct, flowing, Gaussian-shaped features in the imaging acquisition following SVD-filtering (**Figure 1B**). These diffraction-limited microbubble features could be extracted and localized at a sub-diffraction precision and tracked frame-to-frame to produce flow trajectories yielding both the location and velocity of the microvasculature containing the contrast agent. Conventional contrast power processing (**Figure 1C**) demonstrates the blood volume of the mouse brain but lacked the ability to distinguish microvasculature. After accumulating all the microbubble trajectories, the final ULM image (**Figure 1D**) demonstrates well defined microvascular reconstructions for both superficial (cortical) and deep-brain structures (e.g.: hippocampus) and includes flow direction and velocity information which may help distinguish between artery and vein flows for select anatomies. Post-processing of the super-resolution vessel map allowed for velocity profile analysis of individual microvessels, as demonstrated in the hippocampal example show in **Figure 1D**. Generally, as the vessel was traversed longitudinally, we observed that the vessel diameter and peak velocity decreased. The difference between the peak velocity and the mean velocity profiles also reduced, which may imply that there are reduced levels of pulsatility in higher order vessels.

### The 5xFAD cohort demonstrates hypo-perfusion in deep brain structures in comparison to WT control

Super-resolution ULM directional vessel maps and velocity maps were reconstructed for the WT and 5xFAD cohorts in this study, as demonstrated in **Figure 2A-B**. Manual segmentations were performed on these imaging cross-sections with the aid of the Allen brain atlas to produce ROIs corresponding to the dorsal isocortex (region **i**), the entorhinal cortex (region **ii**), and the hippocampal region (region **iii**). Poor vascular reconstruction was noted for the most lateral portion of the cortex, possibly due to interference from strong acoustic scattering from the ear bar, or from acoustic shadowing from the remaining portion of the lateral skull. Quantification of regional anatomy measures of vascularity, median flow velocity, and the sum-of-angles metric, a measure of tortuosity, are demonstrated in **Figure 2C-E**. The vascularity index was a measure of the total percentage of the ROI, which was comprised of reconstructed vasculature, a metric that is roughly analogous to microvascular density. All anatomical regions demonstrated a statistically significant decrease in vascularity for the 6-month 5xFAD animals in comparison to the WT control (isocortex: 5xFAD = 39.1 ± 16.8 % vs. WT = 49.6 ± 13.2 %, **p = 0.005**; entorhinal cortex: 5Xfad = 7.9 ± 6.2 % vs. WT = 14.1 ± 10.9 %, **p = 0.004**, hippocampal region: 5xFAD = 49.2 ± 16.0 % vs. WT = 61.3 ± 11.7 %, **p < 0.001**), with no evidence of a decrease in the younger 3-month-old cohort (isocortex: 5xFAD = 42.6 ± 21.2 % vs. WT = 41.4 ± 23.9 %, p = 0.85; entorhinal cortex: 5xFAD = 8.2 ± 7.4 % vs. WT = 9.4 ± 8.8 %, p = 0.62, hippocampal region: 5xFAD = 39.8 ± 20.2 % vs. WT = 48.5 ± 11.6 %, p = 0.06) (**Figure 2C**).

The 3-month cohort had a significant decrease in flow velocity for the entorhinal cortex and hippocampal regions (isocortex: 5xFAD = 4.62 ± 0.76 mm/s vs. WT = 4.60 ± 0.73 mm/s, p = 0.94; entorhinal cortex: 5xFAD = 3.48 ± 0.32 mm/s vs. WT = 3.99 ± 0.53 mm/s, **p < 0.001**, hippocampal region: 5xFAD = 4.82 ± 0.40 mm/s vs. WT = 5.32 ± 0.56 mm/s, **p < 0.001**), with a sustained decrease in velocity for all ROIs in the 6-month group (isocortex: 5xFAD = 3.77 ± 0.53 mm/s vs. WT = 4.55 ± 0.48 mm/s, **p < 0.001**; entorhinal cortex: 5xFAD = 3.61 ± 0.81 mm/s vs. WT = 4.35 ± 0.44 mm/s, **p < 0.001**, hippocampal region: 5xFAD = 4.97 ± 0.79 mm/s vs. WT = 5.54 ± 0.65 mm/s, **p < 0.001**) (**Figure 2D**). The sum-of-angles tortuosity metric was found to be significantly higher in the 3-month 5xFAD animals in the entorhinal cortex and the hippocampal region (isocortex: 5xFAD = 1.30 ± 0.16 a.u. vs. WT = 1.30 ± 0.27 a.u., p = 0.98; entorhinal cortex: 5xFAD = 1.66 ± 0.52 a.u. vs. WT = 1.28 ± 0.37 a.u., **p = 0.004**, hippocampal region: 5xFAD = 1.24 ± 0.18 a.u. vs. WT = 1.11 ± 0.19 a.u., **p = 0.013**), and was found to be elevated in all regions for the 6-month old 5xFAD animals (isocortex: 5xFAD = 1.75 ± 0.34 a.u. vs. WT = 1.38 ± 0.18 a.u., **p < 0.001**; entorhinal cortex: 5xFAD = 1.92 ± 0.45 a.u. vs. WT = 1.48 ± 0.29 a.u., **p < 0.001**, hippocampal region: 5xFAD = 1.22 ± 0.26 a.u. vs. WT = 1.10 ± 0.18 a.u., **p = 0.027**) (**Figure 2E**). Taken together, these results may imply that there is a functional impairment of vasculature (velocity, tortuosity) in the 5xFAD model which precedes structure changes (vascularity) of the cerebrovascular network.

### Local vessel analysis shows blood velocity reduction in lateral cerebral arteries and in penetrating cortical vessels

As a case study on the investigative power of ULM imaging, individual vessels were selected and analyzed from super-resolution reconstructions from both WT and 5xFAD animals, as demonstrated in local ROIs in **Figure 3**. Velocity profiles were reconstructed for vessels of interest with both the lateral peak velocity estimates and a parabolic profile fit under the assumption of laminar blood flow. In this case example, the lateral cerebral artery of the 5xFAD animal demonstrated a substantial decrease in peak velocity in comparison to the WT control (**Figure 3A**). Regions of cortical vessels were also investigated (**Figure 3B**). The directional flow mapping for the cortical region may give insight into artery vs. vein flow in the cortex. It was found that both the penetrating cortical artery (vessel **iii** and **v**) and penetrating cortical vein (vessel **iv** and **vi**) demonstrated a decrease in peak velocity for the 5xFAD animal. Finally, a hierarchical analysis of a penetrating cortical vessel was performed to confirm that the velocity profiles are consistent with the expected result of decreased velocity as you travel along the length of the vessel (**Figure 3C**). Interestingly, for the smallest diameter vessel sections (denoted as **d** and **g**), the peak velocity and mean velocity estimates were nearly overlapping, which may imply a more steady-state or plug flow in the section of the vessel.

### ULM demonstrates good correspondence to histology with Aβ co-localized with hypo-perfusion

Co-registered ULM images with histological sections are demonstrated in **Figure 4**. It was found that there was a good overall correspondence between the FITC dextran staining on histology (green channel) and the ULM vascular reconstruction images, providing evidence that the ULM imaging is a physiologically relevant representation of the vasculature. The WT animal had a high degree of vascularization throughout the entire brain in both ULM and histology, whereas the 5xFAD animal demonstrated a profound decrease in vasculature for the hippocampal and entorhinal cortex regions in the 5xFAD example. Amyloid beta plaque deposition is demonstrated in the red channel of the 5xFAD animal (**Figure 4B**), with substantial levels of Aβ in the hippocampal and subcortical regions of the brain. The WT control did not show any evidence of Aβ deposition. Regions of high amyloid beta deposition showed a co-localization with reduced microvascular staining on histology and regionalized hypo-perfusion in the ULM images.

**Figure 4.**
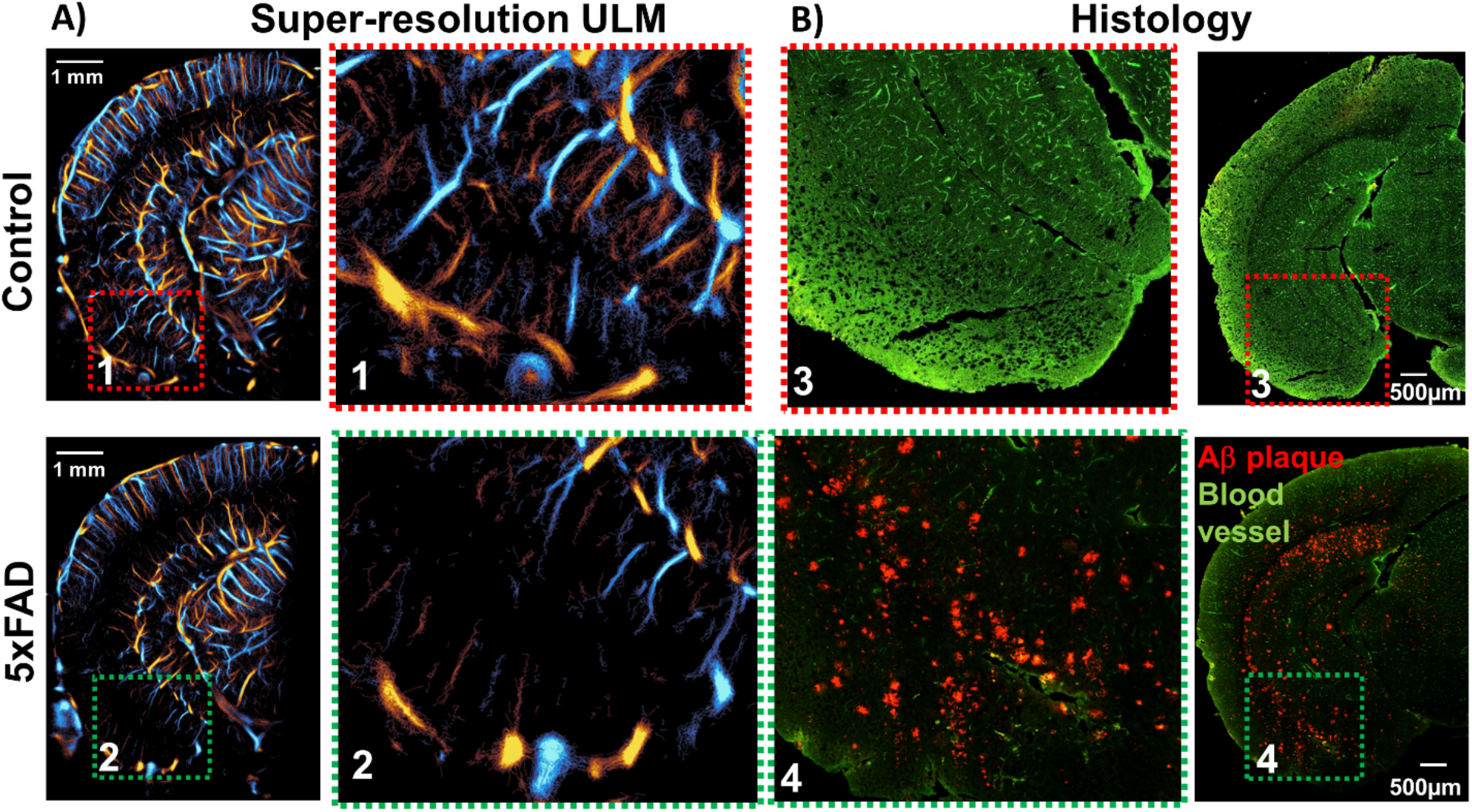
Comparison of ULM to histological analysis of vasculature and Aβ plaques. **(A)** Super-resolution ULM images demonstrate a profound decrease in vasculature for the hippocampal and entorhinal cortex regions in this animal comparison. **(B)** The co-registered histological sections from these animals reveal a good correspondence between ULM vascular reconstruction and histological staining of the microvasculature (green channel; FITC dextran). The regions of high amyloid beta deposition (red channel) showed a co-localization with regions of reduced microvascular staining on histology and reduced presence of microvessels in the ULM images.

## Discussion

This study investigated the vascular pathology present in the 5xFAD mouse model of AD using super-resolution ULM imaging. We demonstrated that this technology provides microvascular-scale characterization of cerebrovascular impairments ranging from superficial to deep-brain structures, most notably for the hippocampus. This allows for anatomy-based regional analysis to observe global shifts in vascular dynamics due to AD progression and for the examination of the correspondence between microvascular structure and Aβ deposition, while also permitting individual vessel characterization to elucidate localized impairments in function. ULM imaging also provides directional flow information that may be able to decouple arteriole and venous vascular features for select anatomical regions (e.g.: dorsal penetrating cortical vessels). A case study was also demonstrated for hierarchical analysis of hippocampal and cortical vessels, which yielded the expected trend of decreasing peak velocity and reduced pulsatility as the vessels were traversed longitudinally.

We found that the 5xFAD mouse model demonstrated an early stage (3-month-old) decrease in hippocampal and entorhinal cortex median blood flow velocity, which progressed into a deficit in vascular density in the 6-month-old animal cohort, along with sustained reductions in blood velocity for these regions (**Fig 2**). We further demonstrated that this decrease in blood flow velocity was not exclusive to the hippocampal or entorhinal cortex regions: a local analysis of lateral cerebral arteries and penetrating cortical vessels revealed a reduced peak velocity in 5xFAD mice relative to WT control (**Fig 3**). Global quantifications from the isocortex also demonstrated a decrease in vascularity and velocity in the 6-month cohort, but not in the younger cohort (**Fig 2**). The sum-of-angles metric, an establish index of vascular tortuosity, was found to be elevated in the 6-month 5xFAD cohort in all regions and elevated in the entorhinal cortex and hippocampal region of the 3-month 5xFAD cohort, which may be indicative of aberrant vascular function and/or perfusion deficits. Taken together, these results may imply that there is a functional impairment of the vasculature at a relatively early time point in the 5xFAD model, which only manifests as structural differences later in the development of the pathology, possibly due to aberrant or pathological vascular remodeling in response to regional stress. These findings can be contextualized with reports that AD associated cognitive deficits in 5xFAD and SwDI animals are correlated to cerebral microvascular amyloid deposition but not to parenchymal amyloid plaques or to total Aβ levels (Xu et al., 2014). This implies that the amyloid-vascular interplay is critical for the development of behavioral deficits in this animal model, however the exact mechanism and manifestation of this interplay is currently unclear. The ULM imaging results suggest that early-stage cerebral microvascular amyloid presents as functional deficits in cerebrovascular flow.

When compared to gold-standard histological analysis of co-registered brain sections we found a good correspondence between vascular FITC staining and ULM maps of cerebral blood flow throughout the entire depth of the brain (**Fig 4**), which provides confidence that ULM is accurately reconstructing patent cerebrovasculature. Notably, Aβ plaque deposition was co-localized with regional hypo-perfusion in histological sections and in corresponding regions of ULM images (**Fig 4**). This is consistent with similar findings in literature that report that cerebral amyloid angiopathy, and the corresponding deficits in vascular perfusion and progression in microvascular damage, occur alongside the accumulation of Aβ in the 5xFAD model (Giannoni et al., 2016).

This study has some limitations which should be discussed to better interpret the results presented. ULM reconstruction of microvasculature is inherently stochastic: vessels are perfused gradually, sparsely, and randomly by MBs through a process that is dictated by physiology and which necessitates long imaging durations to ensure adequate vascular sampling (Christensen-Jeffries et al., 2019; Hingot et al., 2019; Lowerison et al., 2020a). This has implications not only for the pragmatic challenges of the technology (e.g.: difficulty with clinical translation, risk of animal death, obstacle to longitudinal study design), but also introduces several sources of variability into the microvascular quantifications extracted from ULM images. Tissue motion poses a persistent risk to super-resolution ULM reconstruction due to drift in MB localizations, which causes misregistration in the final accumulation. Although motion compensation was performed in the ULM processing pipeline, it in unable to correct for out-of-plane shifts or for tissue deformations (e.g.: tissue motion from large vessel pulsatility). Furthermore, the long imaging duration may impart real physiological changes that can either obscure, or exasperate, baseline differences in AD cerebrovasculature in comparison to WT animals. In this study, mice were anesthetized with vaporized isoflurane for a typical duration of about 2 hours, a duration which impacts cardiac output of the animal and may also induce shifts cerebrovascular perfusion. It has also been reported in literature that isoflurane has a dose-dependent dilatory effect on the cerebral blood flow (Matta et al., 1999), which may have differential consequences on AD mice and WT animals. Another consideration was that the craniotomy was performed just before imaging, which poses the risk of superficial cortical damage. Although care was taken during ULM image analysis to avoid any areas of obvious cortical injury, there may be out-of-plane damage which influenced the final ULM accumulation result. This also introduces the confounding effects of localized inflammation and vascular flow responses to trauma.

ULM imaging is best performed under conditions of ‘ideal’ MB concentration; that is MB signals that are sufficiently spatially sparse to provide unambiguous localization of centroid positions, but with suitable MB numbers to ensure adequate perfusion of microvasculature within the prescribed imaging acquisition time. However, commercial MB contrast agents have a wide distribution of post-activation concentrations, which introduces uncertainty into the perfusion rate of contrast in terms of MBs delivered per unit time. This is compounding by the technical challenge of tail-vein catheterization, which impacts the variability of intravenous delivery of contrast. Although the fixed imaging acquisition duration used in this study (64,000 frames) should be sufficient to reconstruct the majority of cerebrovasculature, there is the possibility that microvascular perfusion did not reach the asymptote of accumulation saturation for particularly slow flowing regions. This particularly implicates the vascularity deficit seen in the 6-month-old 5xFAD animals, which also demonstrated substantially reduced blood flow velocity. Finally, the positioning of the securing ear-bars and the placement of the cranial window yielded strong reflector artifacts in the deep regions of the brain near to the entorhinal cortex. These artifacts may explain the low amount of vascularity that was observed in the entorhinal region relative to the rest of the brain, as the strong signal artifacts obscured MB features in some cases.

We explored the use of ULM imaging for local and hierarchical vessel analysis. However, the uncertainty in the elevational positioning of the coronal section of the brain made it impossible to confirm that we were examining corresponding vessels between 5xFAD and WT animals. This severely limited the conclusions that could be drawn from either the individual or the hierarchical vessel analysis, beyond serving as an internal validation. A similar limitation also applies to the co-registration of histological sections with ULM imaging planes. The elevational beamwidth of ultrasound is substantially larger than the slice thickness of histology, which limits a more refined analysis into the co-localization of vascular hypo-perfusion with Aβ deposition.

We have demonstrated the use of super-resolution ULM on the 5xFAD mouse model of AD (3-month and 6-month-old cohorts), in comparison to age and sex-matched WT controls, to reveal microvascular scale reconstructions throughout the whole brain depth. We found that functional decreases in hippocampal and entorhinal flow velocity preceded structural deficits in regional vascular density. Analysis of local blood profiles in the cortex and in lateral arteries further demonstrated a consistent flow velocity reduction in the 5xFAD group for different brain regions. We also found that flow velocity decreases with increasing vascular tree order. These data suggest that this novel imaging technology can be used to investigate the pathophysiology of AD, and evaluate novel therapeutic approaches that impact the deep microvasculature of mouse models of AD.

## Acknowledgements

This study was partially supported by the National Cancer Institute, the National Institute of Biomedical Imaging and Bioengineering, the National Institute on Deafness and Other Communication Disorders, and the National Institute on Aging of the National Institutes of Health under grant numbers R00CA214523, R21EB030072, R21DC019473 and R03AG059103, as well as a grant to DL from the Kiwanis Neuroscience Research Foundation. The content is solely the responsibility of the authors and does not necessarily represent the official views of the National Institutes of Health. NCS and MRL are supported by Beckman Institute Postdoctoral Fellowships.

## Notes

**Conflict of interest statement:** The authors declare no competing financial interests.

### Competing Interest Statement

The authors have declared no competing interest.

## References

Ahtiluoto S, Polvikoski T, Peltonen M, Solomon A, Tuomilehto J, Winblad B, Sulkava R, Kivipelto M (2010) Diabetes, Alzheimer disease, and vascular dementia: a population-based neuropathologic study. Neurology 75:1195–1202.

Bellew KM, Pigeon JG, Stang PE, Fleischman W, Gardner RM, Baker WW (2004) Hypertension and the rate of cognitive decline in patients with dementia of the Alzheimer type. Alzheimer Dis Assoc Disord 18:208–213.

Bloom GS (2014) Amyloid-β and tau: the trigger and bullet in Alzheimer disease pathogenesis. JAMA Neurol 71:505–508.

Christensen-Jeffries K, Brown J, Harput S, Zhang G, Zhu J, Tang M, Dunsby C, Eckersley RJ (2019) Poisson Statistical Model of Ultrasound Super-Resolution Imaging Acquisition Time. IEEE Transactions on Ultrasonics, Ferroelectrics, and Frequency Control 66:1246–1254.

Christensen-Jeffries K, Browning RJ, Tang M-X, Dunsby C, Eckersley RJ (2015) In vivo acoustic super-resolution and super-resolved velocity mapping using microbubbles. IEEE Trans Med Imaging 34:433–440.

de la Torre JC (1994) Impaired brain microcirculation may trigger Alzheimer’s disease. Neurosci Biobehav Rev 18:397–401.

de la Torre JC, Mussivand T (1993) Can disturbed brain microcirculation cause Alzheimer’s disease? Neurol Res 15:146–153.

Demené C, Robin J, Dizeux A, Heiles B, Pernot M, Tanter M, Perren F (2021) Transcranial ultrafast ultrasound localization microscopy of brain vasculature in patients. Nat Biomed Eng 5:219–228.

Demeulenaere O, Bertolo A, Pezet S, Ialy-Radio N, Osmanski B, Papadacci C, Tanter M, Deffieux T, Pernot M (2022) In vivo whole brain microvascular imaging in mice using transcranial 3D Ultrasound Localization Microscopy. EBioMedicine 79:103995.

Desailly Y, Pierre J, Couture O, Tanter M (2015) Resolution limits of ultrafast ultrasound localization microscopy. Phys Med Biol 60:8723–8740.

Desailly Y, Tissier A-M, Correas J-M, Wintzenrieth F, Tanter M, Couture O (2017) Contrast enhanced ultrasound by real-time spatiotemporal filtering of ultrafast images. Phys Med Biol 62:31–42.

Errico C, Pierre J, Pezet S, Desailly Y, Lenkei Z, Couture O, Tanter M (2015) Ultrafast ultrasound localization microscopy for deep super-resolution vascular imaging. Nature 527:499–502.

Giannoni P, Arango-Lievano M, Neves ID, Rousset M-C, Baranger K, Rivera S, Jeanneteau F, Claeysen S, Marchi N (2016) Cerebrovascular pathology during the progression of experimental Alzheimer’s disease. Neurobiol Dis 88:107–117.

Glenner GG, Wong CW (1984) Alzheimer’s disease: initial report of the purification and characterization of a novel cerebrovascular amyloid protein. Biochem Biophys Res Commun 120:885–890.

Hardy J, Allsop D (1991) Amyloid deposition as the central event in the aetiology of Alzheimer’s disease. Trends Pharmacol Sci 12:383–388.

HHS (2021) National Plan to Address Alzheimer’s Disease: 2021 Update. :117.

Hingot V, Errico C, Heiles B, Rahal L, Tanter M, Couture O (2019) Microvascular flow dictates the compromise between spatial resolution and acquisition time in Ultrasound Localization Microscopy. Scientific Reports 9:2456.

Hobby JD (1986) Smooth, easy to compute interpolating splines. Discrete Comput Geom 1:123–140.

Huang C, Lowerison MR, Trzasko JD, Manduca A, Bresler Y, Tang S, Gong P, Lok U-W, Song P, Chen S (2020) Short Acquisition Time Super-Resolution Ultrasound Microvessel Imaging via Microbubble Separation. Scientific Reports 10:1–13.

Iadecola C (2004) Neurovascular regulation in the normal brain and in Alzheimer’s disease. Nat Rev Neurosci 5:347–360.

Jaqaman K, Loerke D, Mettlen M, Kuwata H, Grinstein S, Schmid SL, Danuser G (2008) Robust single-particle tracking in live-cell time-lapse sequences. Nature Methods 5:695–702.

Karran E, Mercken M, Strooper BD (2011) The amyloid cascade hypothesis for Alzheimer’s disease: an appraisal for the development of therapeutics. Nat Rev Drug Discov 10:698–712.

Kisler K, Nelson AR, Montagne A, Zlokovic BV (2017) Cerebral blood flow regulation and neurovascular dysfunction in Alzheimer disease. Nat Rev Neurosci 18:419–434.

Kivipelto M, Ngandu T, Fratiglioni L, Viitanen M, Kåreholt I, Winblad B, Helkala E-L, Tuomilehto J, Soininen H, Nissinen A (2005) Obesity and vascular risk factors at midlife and the risk of dementia and Alzheimer disease. Arch Neurol 62:1556–1560.

Lowerison MR, Huang C, Kim Y, Lucien F, Chen S, Song P (2020a) In Vivo Confocal Imaging of Fluorescently Labeled Microbubbles: Implications for Ultrasound Localization Microscopy. IEEE Transactions on Ultrasonics, Ferroelectrics, and Frequency Control 67:1811–1819.

Lowerison MR, Huang C, Lucien F, Chen S, Song P (2020b) Ultrasound localization microscopy of renal tumor xenografts in chicken embryo is correlated to hypoxia. Scientific Reports 10:1–13.

Lowerison MR, Sekaran NVC, Zhang W, Dong Z, Chen X, Llano DA, Song P (2022) Aging-related cerebral microvascular changes visualized using ultrasound localization microscopy in the living mouse. Sci Rep 12:619.

Matta BF, Heath KJ, Tipping K, Summors AC (1999) Direct cerebral vasodilatory effects of sevoflurane and isoflurane. Anesthesiology 91:677–680.

Matthews KA, Xu W, Gaglioti AH, Holt JB, Croft JB, Mack D, McGuire LC (2019) Racial and ethnic estimates of Alzheimer’s disease and related dementias in the United States (2015-2060) in adults aged ≥65 years. Alzheimers Dement 15:17–24.

Nortley R, Korte N, Izquierdo P, Hirunpattarasilp C, Mishra A, Jaunmuktane Z, Kyrargyri V, Pfeiffer T, Khennouf L, Madry C, Gong H, Richard-Loendt A, Huang W, Saito T, Saido TC, Brandner S, Sethi H, Attwell D (2019) Amyloid β oligomers constrict human capillaries in Alzheimer’s disease via signaling to pericytes. Science 365:eaav9518.

R Core Team (2019) R: A Language and Environment for Statistical Computing. Vienna, Austria: R Foundation for Statistical Computing. Available at: https://www.R-project.org/.

Schuff N, Woerner N, Boreta L, Kornfield T, Shaw LM, Trojanowski JQ, Thompson PM, Jack CR Jr, Weiner MW, the Alzheimer’s; Disease Neuroimaging Initiative (2009) MRI of hippocampal volume loss in early Alzheimer’s disease in relation to ApoE genotype and biomarkers. Brain 132:1067–1077.

Sert NP du et al. (2020) Reporting animal research: Explanation and elaboration for the ARRIVE guidelines 2.0. PLOS Biology 18:e3000411.

Shelton SE, Lee YZ, Lee M, Cherin E, Foster FS, Aylward SR, Dayton PA (2015) Quantification of microvascular tortuosity during tumor evolution utilizing acoustic angiography. Ultrasound Med Biol 41:1896–1904.

Song P, Manduca A, Trzasko JD, Chen S (2017a) Ultrasound Small Vessel Imaging With Block-Wise Adaptive Local Clutter Filtering. IEEE Transactions on Medical Imaging 36:251–262.

Song P, Manduca A, Trzasko JD, Chen S (2017b) Noise Equalization for Ultrafast Plane Wave Microvessel Imaging. IEEE Trans Ultrason Ferroelectr Freq Control 64:1776–1781.

Song P, Manduca A, Trzasko JD, Daigle RE, Chen S (2018a) On the Effects of Spatial Sampling Quantization in Super-Resolution Ultrasound Microvessel Imaging. IEEE Trans Ultrason Ferroelectr Freq Control 65:2264–2276.

Song P, Trzasko JD, Manduca A, Huang R, Kadirvel R, Kallmes DF, Chen S (2018b) Improved Super-Resolution Ultrasound Microvessel Imaging with Spatiotemporal Nonlocal Means Filtering and Bipartite Graph-Based Microbubble Tracking. IEEE Trans Ultrason Ferroelectr Freq Control 65:149–167.

Tang S, Song P, Trzasko JD, Lowerison M, Huang C, Gong P, Lok U-W, Manduca A, Chen S (2020) Kalman Filter–Based Microbubble Tracking for Robust Super-Resolution Ultrasound Microvessel Imaging. IEEE Transactions on Ultrasonics, Ferroelectrics, and Frequency Control:1–1.

Wickham H (2016) ggplot2: Elegant Graphics for Data Analysis. Springer-Verlag New York. Available at: http://ggplot2.org.

